# Elucidation of SARS-Cov-2 Budding Mechanisms Through Molecular Dynamics Simulations of M and E Protein Complexes

**DOI:** 10.1101/2021.07.26.453874

**Authors:** Logan Thrasher Collins, Tamer Elkholy, Shafat Mubin, David Hill, Ricky Williams, Kayode Ezike, Ankush Singhal

## Abstract

SARS-CoV-2 and other coronaviruses pose major threats to global health, yet computational efforts to understand them have largely overlooked the process of budding, a key part of the coronavirus life cycle. When expressed together, coronavirus M and E proteins are sufficient to facilitate budding into the ER-Golgi intermediate compartment (ERGIC). To help elucidate budding, we ran atomistic molecular dynamics (MD) simulations using the Feig laboratory’s refined structural models of the SARS-CoV-2 M protein dimer and E protein pentamer. Our MD simulations consisted of M protein dimers and E protein pentamers in patches of membrane. By examining where these proteins induced membrane curvature *in silico,* we obtained insights around how the budding process may occur. Multiple M protein dimers acted together to induce global membrane curvature through protein-lipid interactions while E protein pentamers kept the membrane planar. These results could eventually help guide development of antiviral therapeutics which inhibit coronavirus budding.

---

Though much has been learned about the biology of SARS-CoV-2, the part of its life cycle known as budding is still poorly understood. Coronaviruses must bud into the ERGIC in order to form infectious particles.^1^ When expressed together without the help of any other coronavirus proteins, the membrane protein (M protein) and envelope protein (E protein) are sufficient to allow budding of virus-like particles (VLPs) which resemble those produced by wild-type coronaviruses.^2–4^ Yet the exact mechanisms by which the M and E proteins contribute to budding remain unclear. Some have proposed that M proteins oligomerize into a matrix layer to induce membrane curvature,^5,6^ though more recent data on SARS-CoV-2 has indicated that its M proteins might not form such a matrix.^7^ The role of the E protein in budding is also poorly understood, though it is thought to somehow coordinate envelope assembly.^6,8,9^ It should be noted that the M protein is roughly 300 times more abundant in the ERGIC than the E protein.^10^ Expression of the nucleocapsid N protein has also been shown to greatly enhance the yield of budding VLPs compared to when only the M and E protein are present.^11^ By contrast, the famous S protein is not strictly required for coronavirus budding, though it is incorporated into the VLPs when expressed alongside M and E.^3^ Better understanding of budding may open new doors to ways of combating COVID-19.

Molecular dynamics (MD) simulations can help to elucidate biological phenomena, yet there has not been much work involving MD and coronavirus budding. Monje-Galvan and Voth recently performed MD simulations which characterized the movements of individual M protein dimers and individual E protein pentamers in virtual ERGIC membrane.^12^ This revealed some new insights, including that the M protein dimer can introduce local deformations in the membrane. However, their study did not investigate how multiple M dimers or multiple E pentamers might influence membrane curvature, which is important for understanding budding. Yu et al. reported a coarse-grained MD investigation of the completed SARS-CoV-2 virion, which included numerous M, E, and S proteins.^13^ Though the study did involve all of the three structural proteins, it focused on the completed spherical virus rather than on budding. There remains a need for MD simulations of the budding process which interrogate how multiple SARS-CoV-2 structural protein complexes may facilitate budding.

We utilized atomistic MD simulations via GROMACS to investigate the roles of M and E protein complexes in budding. Because of the lack of complete crystal structures of the M and E proteins, we used the Feig laboratory’s predicted structural models of the M protein dimer and E protein pentamer.^14^ We constructed planar membrane patches with lipid composition mimicking that of the ERGIC and inserted transmembrane M and E protein complexes. We ran 800 ns simulations on five systems: a membrane-only system (mem), a system with a single E protein pentamer (1E), a system with four E protein pentamers (4E), a system with a single M protein dimer (1M), and a system with four M protein dimers (4M) (Table S1). Though the focus of our study was on effects from multiple complexes of the same type, we also ran a 400 ns simulation on a system with three M protein dimers and one E protein pentamer (3M1E). One of the most notable outcomes of our simulations was that the 4M system gained a substantial degree of global curvature over time (Fig. 1A), while other systems such as mem had very little curvature (Fig. 1B). To uncover mechanistic insights around these processes, we further performed a series of quantitative analyses on the simulations.

**Figure 1.**
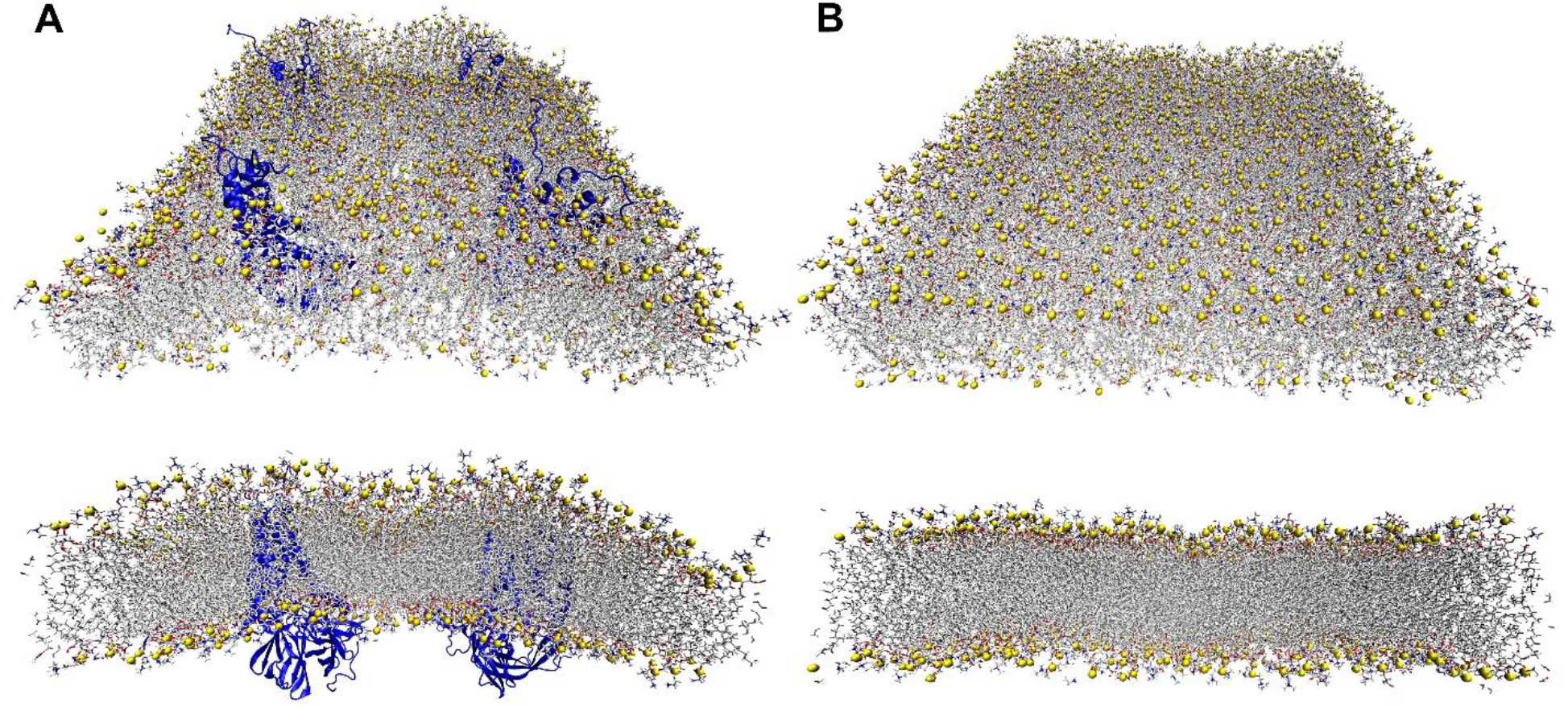
Representative perspective (top) and side-view (bottom) snapshots demonstrating **(A)** strong curvature in the 4M system at 800 ns and **(B)** a lack of substantial curvature in the mem system at 800 ns. To denote the membrane geometry more clearly, phosphorous atoms are shown as yellow spheres. Cytosolic leaflets oriented downwards and lumenal leaflets are oriented upwards.

We first employed g_lomepro^15^ to generate 2D time-averaged mean curvature heatmaps over selected 100 ns intervals (Fig. 2A-E, Fig. S1A) as well as 3D plots of the same data (Fig. 2F-J, Fig. S1B). The 1M system showed a small bulge which grew more pronounced over time, indicating that even lone M protein dimers might induce kinks in the membrane. The 4M system showed by far the highest levels of curvature. Remarkably, the 4M system’s curvature grew both in magnitude and in orderliness over time. In 4M’s 700-800 ns interval, the membrane took the shape of a cylindrical hill, demonstrating the ability of the M proteins to work in an organized fashion. Only small amounts of curvature were visible in the 1E, 4E, and mem simulations, indicating that E protein pentamers may play a role during budding which does not directly involve the induction of curvature. The 3M1E system showed moderate curvature, which was less pronounced than in the 4M system. In summary, these data indicate that E proteins likely do not induce substantial curvature, that isolated M proteins create bulges in the membrane, and that many M proteins together can act together to induce larger amounts of curvature.

**Figure 2.**
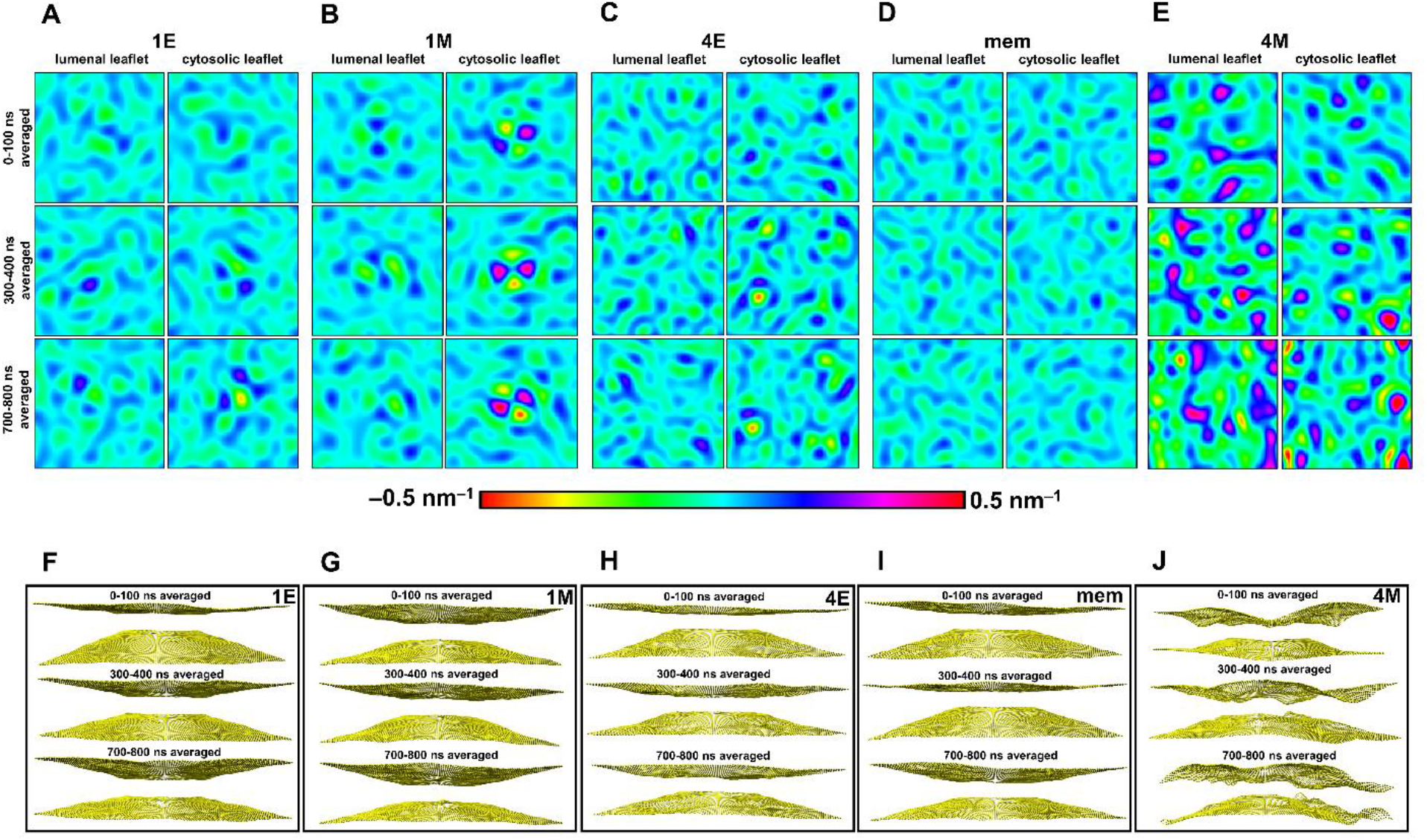
Time-averaged mean curvature heatmaps over selected time intervals for the **(A)** 1E system, **(B)** 1M system, **(C)** 4E system, **(D)** mem system, and **(E)** 4M system and corresponding mean curvature 3D plots for the **(F)** 1E system, **(G)** 1M system, **(H)** 4E system, **(I)** mem system, and **(J)** 4M system. The 3D plots are oriented such that the cytosolic leaflets are oriented downwards and the lumenal leaflets are oriented upwards. All 3D plots are represented as side views of the membranes.

We characterized protein dynamics using MDanalysis^16^ to perform root-mean-square deviation (RMSD) (Fig. 3A-D, Fig. S2A) and radius of gyration (Rg) (Fig. 3E-H, Fig. S2B) calculations and using the GROMACS command line to perform root-mean-square fluctuation (RMSF) calculations (Fig. S3A-D, Fig. S4). By comparison to the M proteins, the E proteins consistently reached higher RMSD values. This is likely due to the unstructured hinge regions connecting the E protein cytosolic α-helices to their transmembrane α-helices, which allowed for more configurational freedom of the cytosolic α-helices. The comparative lack of variability in the M proteins may facilitate retention of their wedgelike shape, which could help induce membrane curvature. Similarly, the Rg values of the M proteins remained relatively constant over time while the Rg values of the E proteins exhibited greater variability over time. RMSF values of M proteins were often high at the residues corresponding to the N and C-terminal unstructured loops, but otherwise remained relatively small in magnitude, supporting the notion that the wedgelike configurations were fairly stiff. RMSF values of E proteins frequently increased around their C-terminal unstructured loops. Though the cytosolic E protein α-helices exhibited high configurational freedom, we observed in VMD that they often adsorbed to each other, resulting in random agglomerations of α-helices (Fig. S5). This could explain why some of the RMSF plots do not show high values around these cytosolic α-helices. The RMSD, Rg, and RMSF data support the notion that the M protein dimers have relatively rigid conformations while the E protein pentamers may have more variable structures.

**Figure 3.**
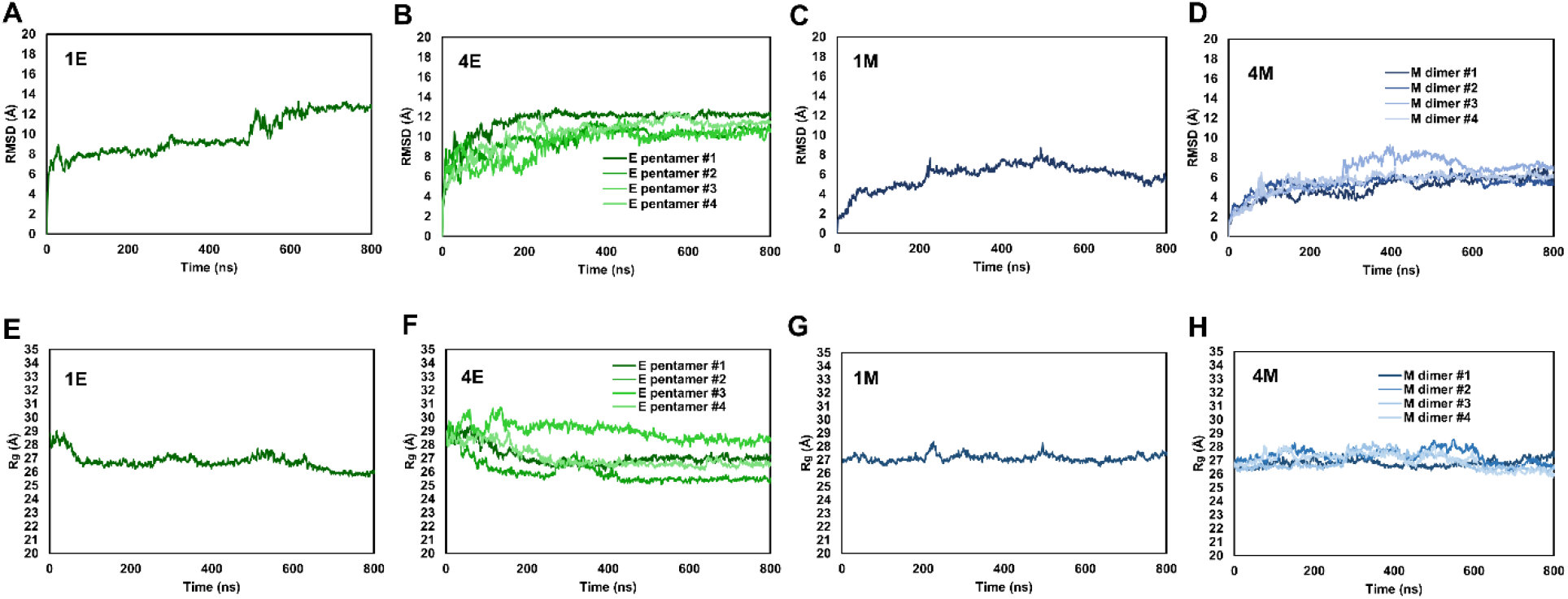
RMSD plots for the **(A)** 1E simulation, **(B)** 4E simulation, **(C)** 1M simulation, and **(D)** 4M simulation as well as Rg plots for the **(E)** 1E simulation, **(F)** 4E simulation, **(G)** 1M simulation, and **(H)** 4M simulation.

To better understand why the M proteins in the 4M system induced such strong curvature, we started by performing time-dependent protein-protein contact analyses on the 4M and 4E simulations to determine whether direct protein-protein interactions were driving the curvature. During the course of the 800 ns trajectory, minimal protein-protein contacts were made between any given pair of M protein dimers (Fig. 4A). The only time that any pair of M dimers came near each other was an isolated incident in the middle of the simulation in which unstructured loops of one of the pairs of M dimers interacted. Since this only occurred between one pair of dimers and only lasted for approximately 100 ns, the event is unlikely to have any functional significance. Furthermore, stochastic protein-protein interactions also happened in the 4E system. As such, the curvature in 4M likely arose from protein-lipid interactions rather than direct protein-protein interactions.

**Figure 4.**
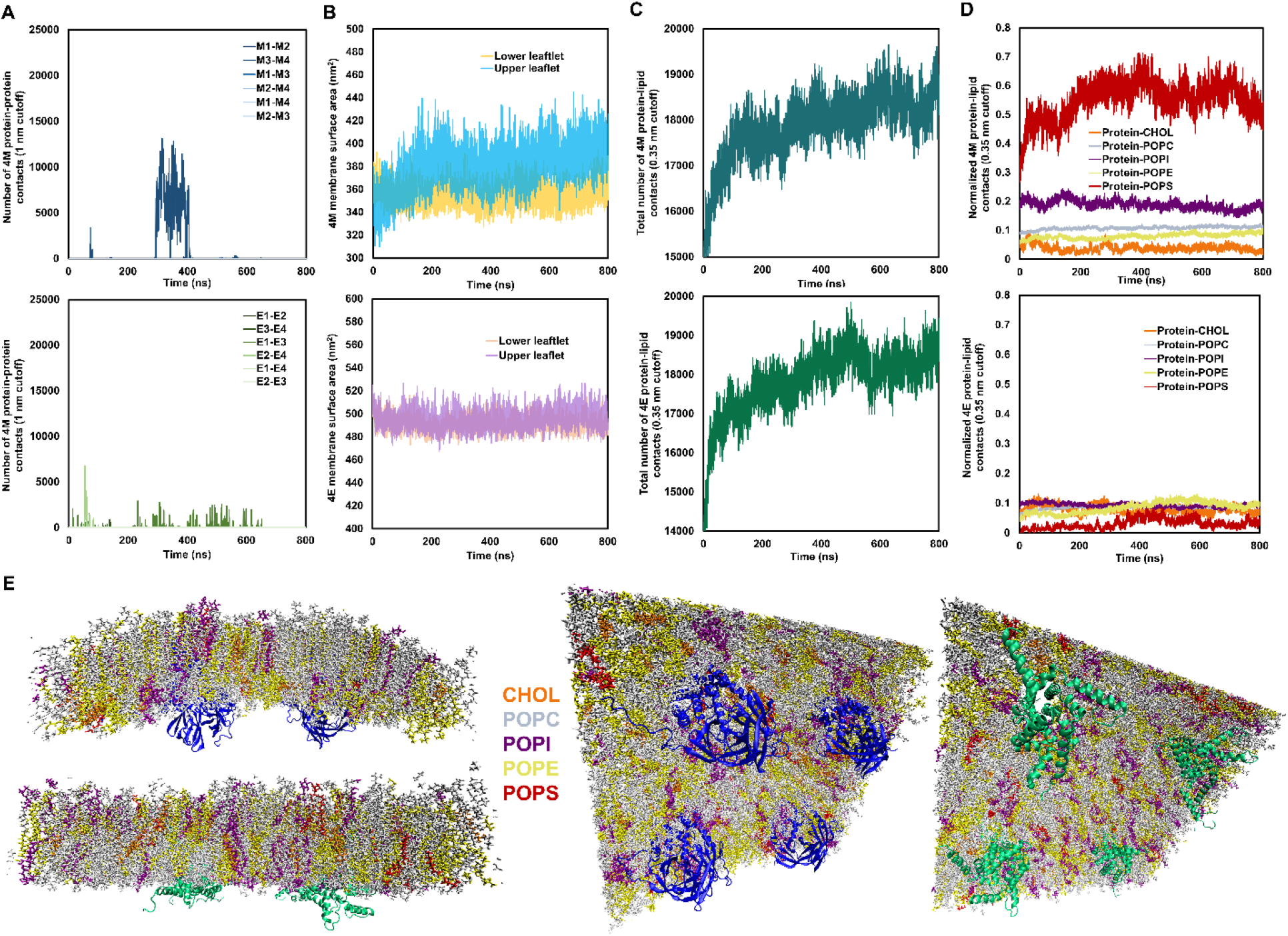
An in-depth comparison of 4M and 4E systems was performed to better understand why 4M showed substantially more curvature. **(A)** Protein-protein contacts among pairs of M dimer complexes in 4M and among pairs of E pentamer complexes in 4E. **(B)** Surface areas of lower membrane leaflets and upper membrane leaflets in 4M and 4E. **(C)** Total protein-lipid contacts in 4M and 4E. **(D)** Normalized protein-lipid contacts by lipid type in 4M and 4E. Normalization was accomplished by dividing the number of contacts for each type of lipid by the number of atoms from that type of lipid in the given system. **(E)** Snapshots of 4M and 4E at 800 ns with lipids colored according to type. CHOL is cholesterol, POPC is phosphatidylcholine, POPI is phosphatidylinositol, POPE is phosphatidylethanolamine, and POPS is phosphatidylserine.

To characterize and compare the protein-lipid dynamics of 4M and 4E, we performed time-dependent membrane surface area calculations (Fig. 4B) and time-dependent protein-lipid contact analyses (Fig. 4C-E). In 4M, the upper membrane leaflet surface area increased over time while the lower membrane leaflet surface area decreased (Fig. 4B). Interestingly, the 4M upper leaflet surface area increased most rapidly during the first 200 ns of the simulation, which was earlier than the period of most dramatic curvature. So, remodeling of the leaflets may have taken place (also see Fig. 2J) in order to achieve the most clearly organized global curvature. We next examined the frequencies of protein-lipid contacts. Total protein-lipid contact frequencies evolved similarly during both the 4M and 4E simulations (Fig. 4C), so we reasoned that the 4M curvature did not come from the growth of the total number of protein-lipid contacts over time. We then computed normalized frequencies of protein-lipid contacts by type of lipid (Fig. 4D). During the 4M simulation, the M dimers displayed substantially greater normalized frequencies of contacts with POPI and POPS lipids compared to CHOL, POPC, and POPE. In the 4E simulation, CHOL, POPC, POPI, and POPE came in contact with the E pentamers at roughly equivalent frequencies, while POPS lipids showed low interaction frequencies. Static snapshots of 4M and 4E at 800 ns support the idea that different types of lipids are distributed in distinct ways between the two systems (Fig. 4E). These data support the notion that the M protein dimers induce curvature through protein-lipid interaction mechanisms rather than by protein-protein interactions as has been hypothesized in the past.^5,6^

Our atomistic MD simulations uncovered insights around the roles of M and E proteins in SARS-CoV-2 budding. Multiple M protein dimers together induced global membrane curvature. Because coronaviruses are known to produce large numbers of M proteins in the ERGIC membranes of infected cells,^10^ we hypothesize that this effect could increase further in the biological reality, leading to enough curvature to encapsulate the RNA genome of the virus. Strikingly, we found that protein-protein interactions did not contribute to the 4M system’s membrane curvature. We instead demonstrated that M dimers remodeled the ERGIC membrane through protein-lipid interactions. RMSD, Rg, and RMSF analyses quantify how the M protein dimers steadily retained their wedge-shaped configuration, indicating that this geometry may have helped sculpt the membrane. Protein-lipid contact analyses demonstrate that M protein dimers preferentially associate with POPS and POPI lipids, suggesting that the M proteins may dynamically reconfigure the ERGIC membrane to create an optimum lipid environment for curvature to occur. POPI lipids specifically are known to facilitate membrane curvature even at low concentrations,^17,18^ so their affinity for the M protein dimers could play a role in stabilizing the membrane in the curved state. Our data indicate that M protein dimers may utilize their wedgelike geometry to mechanically reshape the membrane as well as that the M protein dimers may spatially manipulate POPI and POPS to optimize the creation of a curved membrane.

Lack of curvature in the 1E and 4E simulations indicates that the E protein likely does not directly facilitate membrane curvature during SARS-CoV-2 budding. But since experimental results show that E proteins are essential for budding in coronaviruses,^2–4^ the E protein likely still plays another role in the process. One possibility is that the E protein introduces a planar region into the membrane’s overall curvature profile, eventually creating a viral envelope with a larger radius of curvature than would be possible with only the M proteins. Another possibility is that the E protein orients M proteins with fivefold symmetry to guide them towards inducing spherical curvature for budding. As such, E protein pentamers may organize the behavior of M protein dimers on larger spatiotemporal scales than were possible in our simulations.

Based on the results of our models, we propose that the M protein dimer may represent a valuable target for drugs intended to treat COVID-19 and other coronavirus diseases. To support the idea that the M protein could represent a useful drug target, we submitted the Feig laboratory’s M protein dimer structure to a web server tool called PockDrug.^19^ This tool successfully identified several high-scoring drug pockets (Fig. S6). Due to the high level of conservation of the M protein across different types of coronaviruses,^20^ we postulate that drugs affecting the M protein might have a broad degree of efficacy. Pharmaceuticals which target the M protein could provide a powerful approach by which to mitigate the effects of coronavirus infections.

## Computational Methods

Six MD simulations of M and E proteins in lipid membrane were used in this study. All of the simulations were carried out at atomic resolution using GROMACS 2019.4.^21^ Structures and trajectories were visualized using VMD 1.9.3.^22^ Structures of the E protein pentamer and M protein dimer were obtained from the Feig laboratory’s predicted models.^14^ Six initial configurations were constructed: a membrane-only system (mem), a system with a single E protein pentamer in membrane (1E), a system with four E protein pentamers in membrane (4E), a system with a single M protein dimer in membrane (1M), a system with four M protein dimers in membrane (4M), and a system with three M protein dimers and one E protein pentamer in membrane (3M1E). To mimic the biological ERGIC, the membrane composition used for all six systems was as follows: 57% POPC, 25% POPE, 10% POPI, 2% POPS, 6% CHOL.^14^ All the systems were solvated using explicit water molecules and the appropriate number of potassium counterions was added to each system to prevent long-range electrostatic effects. The CHARMM36 force field^23^ was used for all lipids, ions, and proteins, while the TIP3P^24^ model was implemented for the water molecules. All hydrogen atoms were constrained with the LINCS algorithm,^25^ and long-range electrostatics were evaluated with particle-mesh Ewald summation.^26^ All simulations used 2 fs time step with Leap-Frog integrator^27^ and a 1.4 nm cutoff for all of the interactions. A standard energy minimization procedure was performed using the steepest descent method.^28^ A small NPT equilibration run was performed for each simulation, followed by a production run using a Nose-Hoover thermostat^29^ at 300K and a semi-isotropic pressure coupling with Parinello-Rahmann barostat^30^ at 1 atm. The lengths of the production runs were as follows: 800 ns for mem, 1E, 4E, 1M, and 4M and 400 ns for 3M1E.

Analyses of the results of the simulations included RMSD, Rg, RMSF, time-averaged mean curvature of the membranes, contact analysis among the M proteins in the 4M system, contact analysis between the M proteins and the lipids in the 4M system, and time-dependent membrane surface area calculations for the 4M system. MDanalysis 1.1.1^16^ was used to calculate RMSD and Rg while g_lomepro^15^ was used for the membrane curvature calculations. Each protein’s RMSD was calculated at 0.1 ns intervals by comparing its conformation at a given time step to a reference conformation consisting of the initial equilibrated structure. To correct for the effects of proteins undergoing translations and rotations during the simulation runs, RMSD was adjusted by translating with a vector ***δ*** and rotating with a matrix **R**. In this way, only the changes in the proteins relative to their initial reference structures were included in the final RMSD outputs. The RMSD was calculated using the coordinates of all of the α-carbon atoms in the given protein where **x** describes the coordinates in the current conformation, x_ref_ are the coordinates of the reference conformation, and *n* is the number of α-carbon atoms in the protein.

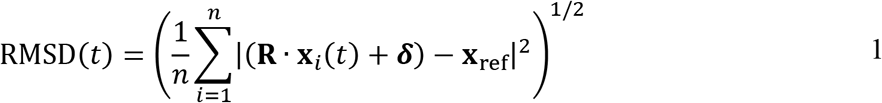

Similarly, R_g_ was calculated for the α-carbon atoms of each protein at 0.1 ns intervals to analyze changes in the compactness of the proteins. R_g_ was computed using the displacement vector **r** between a given protein’s center of mass and each α-carbon of that protein. These calculations were weighted by the mass *m* of the atom in question.

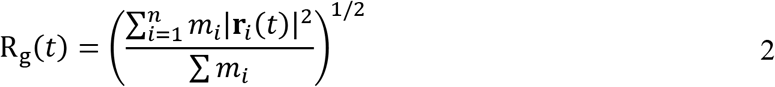

RMSF was calculated using the GROMACS command line for the α-carbon atoms of each protein over the 300-400 ns and 700-800 ns intervals of the simulations. To account for translations and rotations, reference positions from the initial frame of each simulation were included in the commands. GROMACS calculated RMSF at each protein residue *i* using the following equation where *t_j_* describes the series of frames over which the RMSF was computed.

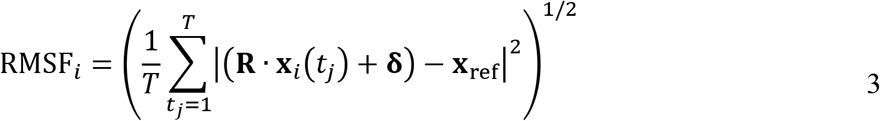

For membrane curvature calculations, the g_lomepro^15^ software package was used to calculate mean curvature as averaged over the frames of the 0-100 ns, 300-400 ns, and 700-800 ns time periods. To analyze the time-dependent frequencies of contacts among the four M protein dimers in the 4M system, the GROMACS hbond command was employed (after appropriate centering). The hbond command was also employed to analyze the time-dependent frequencies of contacts between the four M protein dimers and the lipids in the 4M system. Normalization of the frequencies of contacts among distinct types of lipids was achieved by dividing each frequency value by the number of atoms of the given lipid type in the system under consideration. The FATSLiM software tool was utilized to calculate time-dependent membrane surface area in the 4M system. Performing these quantitative analyses helped us to decipher insights from our simulations.

The PockDrug tool was used to identify predicted drug binding pockets in the M protein dimer.^19^ First, the M dimer structural model from the Feig laboratory^14^ was inputted into the PockDrug web server. The fpocket estimation method was chosen. Five of the top-scoring predicted pockets were selected for visualization in VMD. This technique demonstrated that the M protein dimer may have potential as a drug target.

## Supporting information

Supplementary figures and tables.

## Acknowledgements

Time and resources on the Frontera supercomputer were awarded to Conduit through the COVID-19 High-Performance Computing Consortium project MCB200139. This work used the Extreme Science and Engineering Discovery Environment (XSEDE), which is supported by National Science Foundation grant number ACI-1548562. We thank Ryan Robinson for creating Conduit and bringing the team together.

