## Supporting information for "Elucidation of SARS-Cov-2 Budding Mechanisms Through Molecular Dynamics Simulations of M and E Protein Complexes"

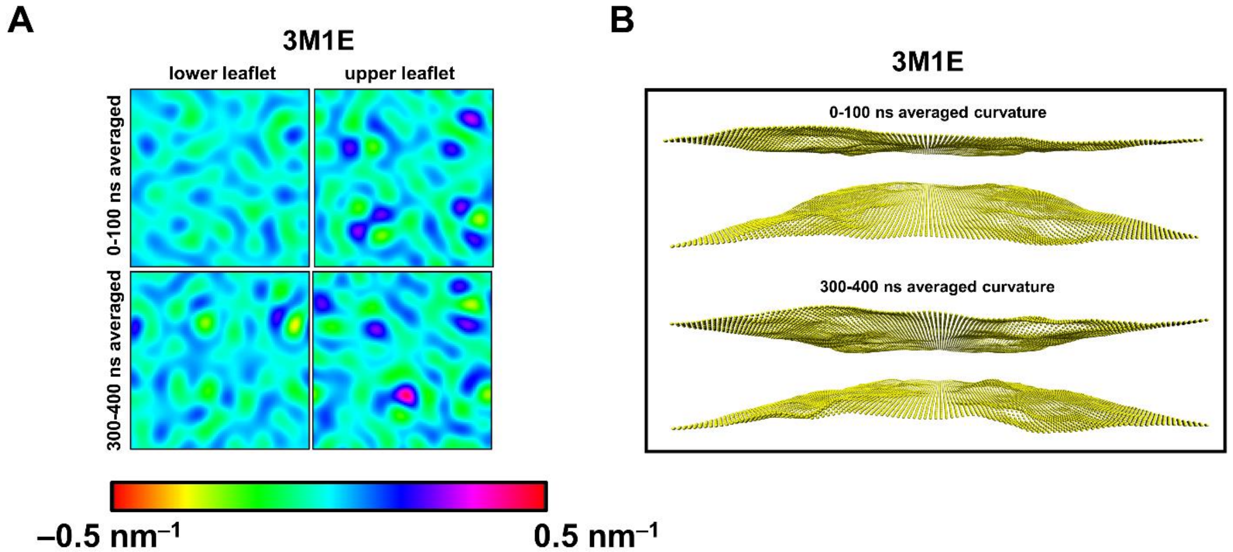


**Supplementary Figure 1 (A)** Time-averaged mean curvature heatmap over selected time intervals for the 3M1E simulation. The lower leaflets are on the cytosolic side of the membrane and the upper leaflets are on the lumenal side of the membrane. **(B)** Time-averaged mean curvature 3D plots over selected time intervals for the 3M1E simulation. The 3D plots are oriented such that the lumenal leaflet is at the top and the cytosolic leaflet is at the bottom.

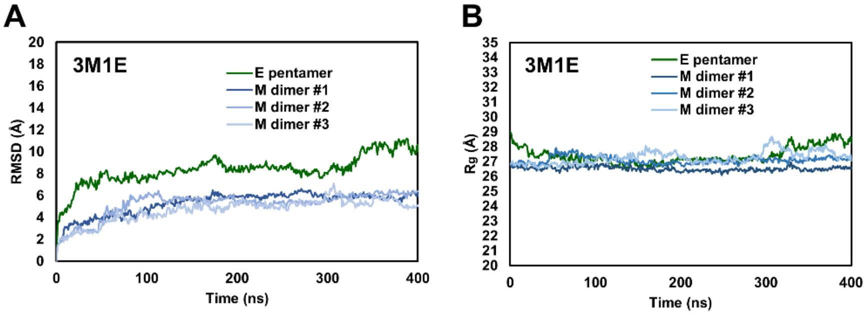


**Supplementary Figure 2 (A)** RMSD plot for the 3M1E simulation and **(B)** $R_g$ plot for the 3M1E simulation.

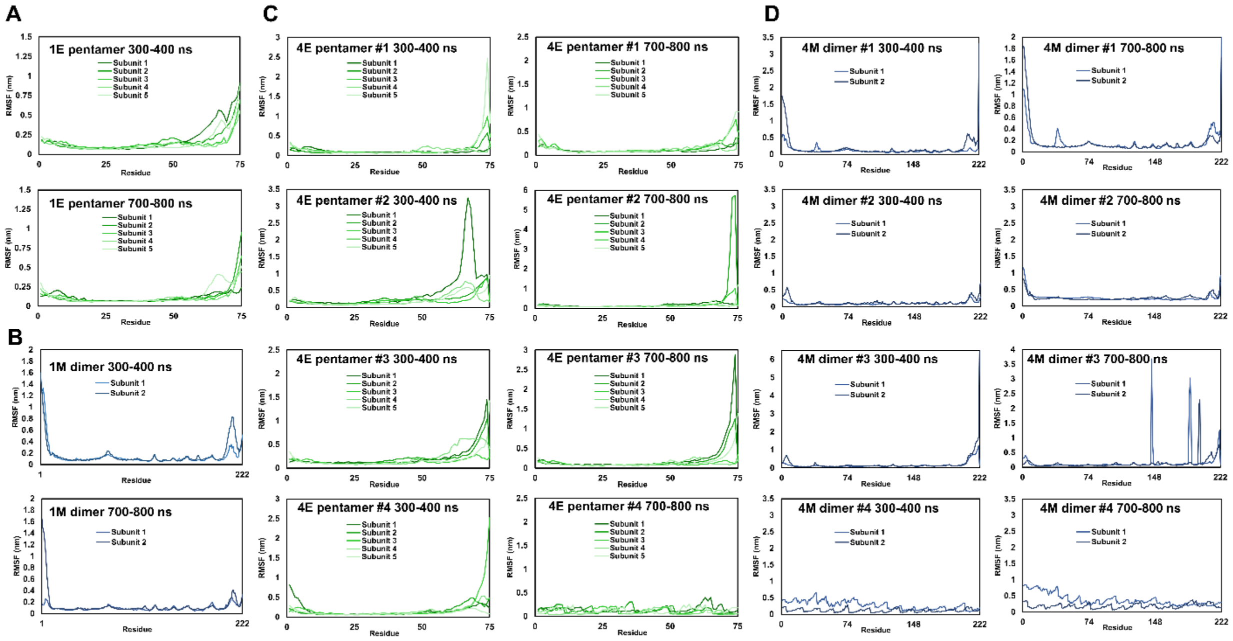


**Supplementary Figure 3** RMSF plots over 300-400 ns and 700-800 ns intervals for each of the proteins in the **(A)** 1E system, **(B)** 1M system, **(C)** 4E system, and **(D)** 4M system.

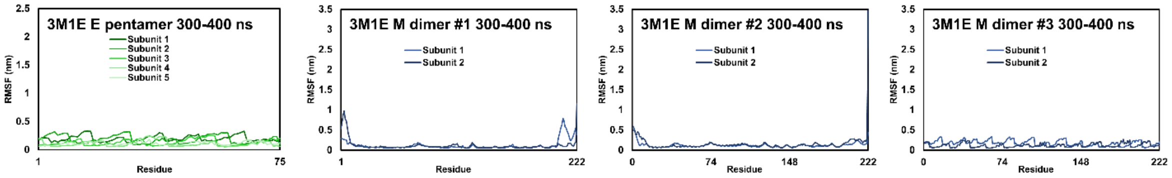


**Supplementary Figure 4** RMSF plots over the 300-400 ns interval for each of the proteins in the 3M1E simulation.

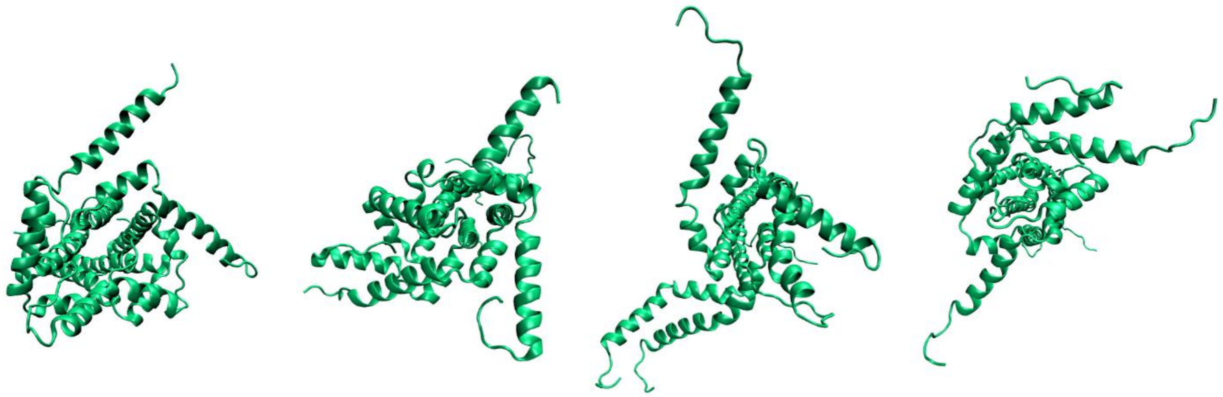

**Supplementary Figure 5** Snapshots of variable E protein configurations arising from the free rotations and random agglomerations of E protein cytosolic α-helices. These snapshots were taken from the 4E system at 800 ns. These E protein pentamers are oriented with their cytosolic domains towards the viewer.

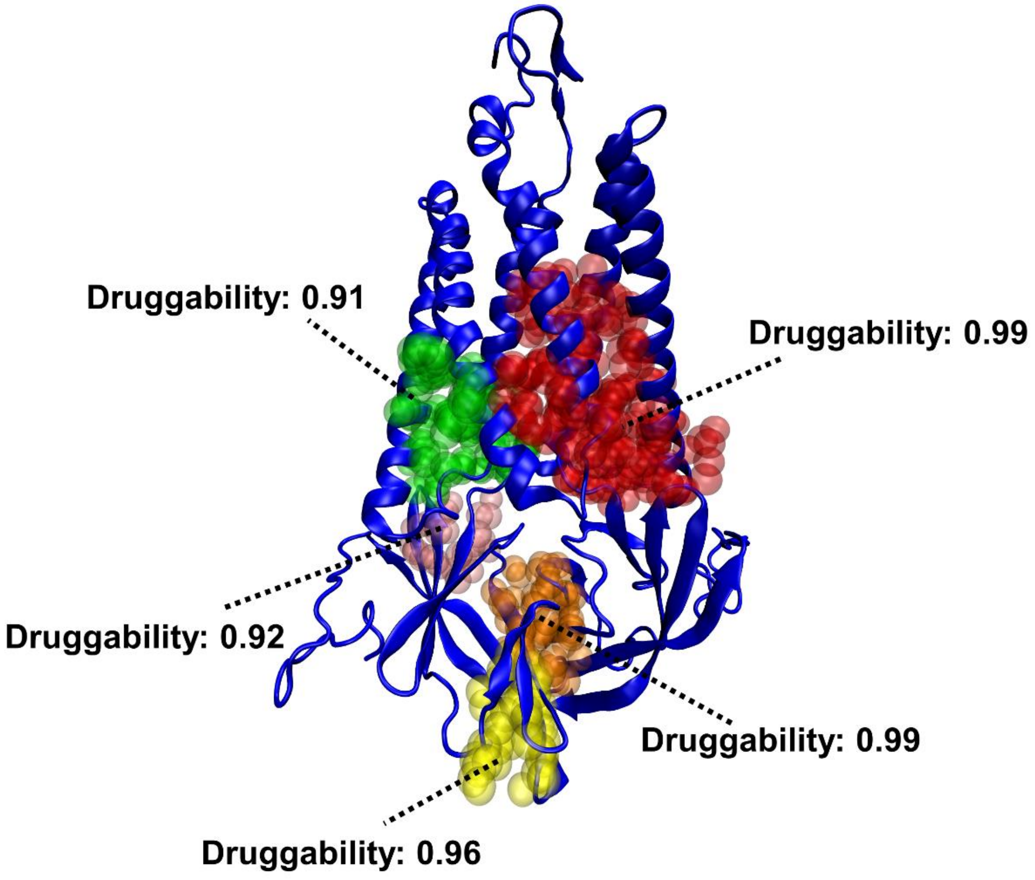


**Supplementary Figure 6** Locations and druggability scores of top drug pockets in the M protein dimer as predicted by the PockDrug tool. While druggability scores of 0.5 or greater correspond to predicted druggable pockets, only the predicted pockets with scores of 0.9 or greater are shown here.

**Supplementary table 1** Sizes of the simulated systems.

| System | Number of atoms |
|---|---|
| 1E | 493115 |
| 1M | 568239 |
| 4E | 607340 |
| 4M | 599868 |
| mem | 405600 |
| 3M1E | 878195 |
